# Controlling brain state prior to stimulation of parietal cortex prevents deterioration of sustained attention

**DOI:** 10.1101/2020.01.06.896357

**Authors:** Grace Edwards, Federica Contò, Loryn K. Bucci, Lorella Battelli

## Abstract

Sustained attention is a limited resource which declines during daily tasks. Such decay is exacerbated in clinical and aging populations. Recent research has demonstrated that inhibition of the intraparietal sulcus (IPS) using low-frequency transcranial magnetic stimulation (LF-rTMS) can lead to an upregulation of functional communication within the attention network. Attributed to functional compensation for the inhibited node, this boost outlasts the stimulation for tens of minutes. Despite the neural change, no behavioral correlate has been found in healthy subjects, a necessary direct evidence of functional compensation. To understand the functional significance of neuromodulatory induced fluctuations on attention, we sought to boost the impact of LF-rTMS through controlling neural excitability prior to LF-rTMS, with the goal to impact behavior. Brain state was controlled using high-frequency transcranial random noise stimulation (HF-tRNS), shown to increase and stabilizes neuronal excitability. Using fMRI-guided stimulation protocols combining HF-tRNS and LF-rTMS, we tested the post-stimulation impact on sustained attention via a multiple object tracking task (MOT). Whilst attention deteriorated across time in the control conditions, HF-tRNS followed by LF-rTMS maintained attention performance up to 94 minutes, doubling the length of successful sustained attention. Multimethod stimulation was also more effective when targeting right IPS, supporting the notion of specialized attention processing in the right hemisphere. Used in a cognitive domain dependent on network-wide neural activity, this tool may be effective in causing lasting neural compensation important for clinical rehabilitation.

## Introduction

Sustained attention is fundamental for cognitively interacting with the environment (DeGangi & Porges, 1990), however, it progressively deteriorates over time (Sarter et al., 2001; Berardi et al., 2001; Whitehurst et al., 2019). This deterioration increases with age (Berardi et al., 2001) and in cognitive disorders, such as ADHD and bipolar disorder (Barkley, 1997; Clark et al., 2005). Thus, a protocol that stabilizes and improves sustained attention for prolonged durations has population-wide application.

Non-invasive brain stimulation can significantly boost cognitive function (Herpich et al. 2019; Reinhart & Nguyen, 2019; Freedberg et al., 2019; Hermiller et al., 2019). Favorable behavioral changes following stimulation may be due to lasting network-wide fluctuation of regions functionally connected to stimulation site (Battelli et al., 2017; Freedberg et al., 2019; Hermiller et al., 2019). In visual attention, low frequency repetitive transcranial magnetic stimulation (LF-rTMS) to the intraparietal sulcus (IPS) is often associated with inhibitory impact on the underlying cortex, and an acute decrease in contralateral attention (Battelli et al., 2009; Edwards et al., 2017). However, following the initial inhibitory effect, LF-rTMS to IPS also results in a cascade of network-wide effects (Capotosto et al., 2012; Szczepanski & Kastner, 2013; Plow et al., 2014; Petitet et al., 2015; Capotosto et al., 2017). For example, inhibition of the IPS with LF-rTMS causes increased functional communication between nodes of the dorsal attention network, 48 minutes post-stimulation (Battelli et al., 2017). One hypothesis is that network-wide functional change compensates for the inhibited node following LF-rTMS (Paus et al., 1997; Grefkes et al., 2009; Lee & D’Esposito, 2012; Plow et al., 2014; Battelli et al., 2017). Functional compensation should predict behavioral benefit lasting the duration of the functional change (Grefkes et al., 2009). Lee & D’Esposito (2012) found theta burst TMS to prefrontal cortex (PFC) increased connectivity in the unstimulated PFC homologue which correlated with a reduced disruption of working memory. Yet such behavioral correlate in attention has not been found following network-wide lasting functional changes after LF-rTMS to IPS in healthy participants (Plow et al., 2014; Battelli et al., 2017). Enduring behavioral change is crucial for non-invasive brain stimulation to be considered in clinical intervention (Edwards et al., 2019).

To affect behavior, we employed a method for boosting the impact of LF-rTMS, and therefore, the following compensatory neuromodulation. Multi-method brain stimulation has been successfully applied to the motor cortex to boost the impact of LF-rTMS, inhibiting physiological response up to an hour after stimulation (Iyer et al., 2003). The lasting inhibition was achieved by applying high-frequency prior to low-frequency stimulation, with the aim of extending the long-term depressive effect (Stanton & Sejnowski, 1989; Christie & Abraham, 1992; Iyer et al., 2003). This effect has been termed “metaplasticity”, the influence of neuronal activation history on subsequent neuronal activity (Abraham & Bear, 1996). To date, multi-method stimulation protocols using a variety of high-frequency and low-frequency stimulation combinations have produced enduring cortical inhibition. These methods have been tested physiologically in the motor cortex using HF-rTMS followed by LF-rTMS (Iyer et al., 2003), anodal transcranial direct current stimulation (a-tDCS) followed by LF-rTMS (Siebner et al., 2004; Bocci et al., 2014), and in the visual cortex using a-tDCS followed by cathodal-tDCS (Fricke et al., 2010), but never in the parietal cortex using a cognitive task as the outcome measure. In order to harness the hypothesized behavioral correlate from functional compensation following LF-rTMS to IPS, we aimed to boost the impact of LF-rTMS over the IPS.

Here, we adapted the multi-method approach to modulate the IPS, a central node of the dorsal attention network, and recorded sustained attention for 94 minutes post-stimulation. In the first experiment, we primed bilateral IPS with high-frequency transcranial random noise stimulation (HF-tRNS), followed by LF-rTMS to initiate long-term inhibition (Christie & Abraham, 1992). We selected HF-tRNS as it has been regularly shown to increase cortical excitability (Terney et al., 2008; Moliadze et al., 2012; Herpich et al., 2018), whilst being well-tolerated by participants (Antal et al., 2017). In the second experiment, we aimed to replicate experiment one, and determine if targeting left or right IPS with multi-method stimulation differentially modulated sustained attention. Furthermore, we ensure our stimulation site selection using fMRI localized left and right IPS, and examined if LF-rTMS alone (without prior HF-tRNS priming) could account for attention change across time.

We hypothesized that sustained attention would decrease with time in the control conditions, whereas, we expected functional compensation following multi-method stimulation would result in improved sustained attention. We conjecture multi-method stimulation would result in sustained inhibition of the IPS, similar to the findings in the motor and visual cortex (Iyer et al., 2003; Bocci et al., 2014) and functional compensation from other nodes of the attention network in response (Paus et al., 1997; Grefkes et al., 2009; Lee & D’Esposito, 2012; Plow et al., 2014; Battelli et al., 2017). In the second experiment, we hypothesized the functional asymmetry previously reported between right and left parietal regions would be evident in post-stimulation behavior in right and left visual fields. The left parietal regions have been shown to orient attention towards the right visual field (Kinsbourne 1977; Kim et al., 1999), therefore left-targeted IPS multi-method stimulation may impact the right visual field only. Whereas the right represents attention in left and right visual space (Corbetta et al., 2005; Battelli et al., 2001; Sheremata et al., 2010; Sheremata & Silver 2015; Ro & Beauchamp, 2020), indicating multi-method stimulation to right IPS should impact both left and right visual fields. Finally, we expected LF-rTMS alone to only initially decrease contralateral visual field attention and have no impact on ipsilateral attention (Dambeck et al., 2006; Battelli et al., 2009).

## Methods & Materials

### Participants

Forty-one volunteers (nineteen females; age range 20-40 years), participated in the two experiments. Each experiment was performed with a new group of participants. All participants gave written informed consent, and the study was approved by Harvard University’s Institutional Review Board: The Human Research Protection Program. In the first experiment, all twenty participants were included in data analysis. In the second experiment, five participants’ data was not analyzed due to incomplete attendance to experiment sessions, leaving sixteen participants in total. All participants had normal to corrected-to-normal vision.

### Experiment 1: Multimethod stimulation to prevent attention deterioration

Experiment 1 was a one-session between-subjects design, whereby participants were randomly assigned to an experimental group when scheduled for the experiment.

#### Stimuli

For behavioral testing during and after stimulation, participants viewed the stimulus on a 13-inch MacBook Pro at a distance of 60 cm (Retina; screen resolution: 1280 × 800). All stimuli were presented using the psychophysical toolbox PsychoPy2 (Peirce et al., 2019).

#### Standard bilateral Multiple Object Tracking Paradigm (MOT)

Participants performed a multiple object tracking paradigm (MOT) for experiments 1 and 2. MOT is a well-established paradigm for recruiting bilateral attention (Pylyshyn & Storm, 1988), and functionally activating bilateral parietal cortices (Culham et al.,1998). On each trial, participants were presented with four objects (black dots, radius 0.25°) either side of a central fixation (Figure 1a). Two objects on either side of the fixation cross flashed (2 Hz for 2 sec) to cue the participant to track these objects amongst the other identical distractors. All objects then moved at a constant speed (degrees per second) for three seconds within a 6° × 6° region and centered 2° to the left and right of the fixation. Speed was set according to the individual participants’ threshold (see *Thresholding* section). Each object repelled one another to maintain a minimum space of 1.5°, never crossed the midline and bounced off of invisible edges within each visual field. After three seconds, the objects stopped moving, one object was highlighted in red, and the participant was asked to respond with a key press to indicate if the highlighted object was the “target” or “distractor” to the objects flashed at the beginning. Importantly, the participant was unaware of which visual field would be tested, necessitating central fixation and tracking within both visual fields simultaneously. After each trial, the fixation point changed to red to indicate incorrect, or green to indicate correct response.

**Figure 1:**
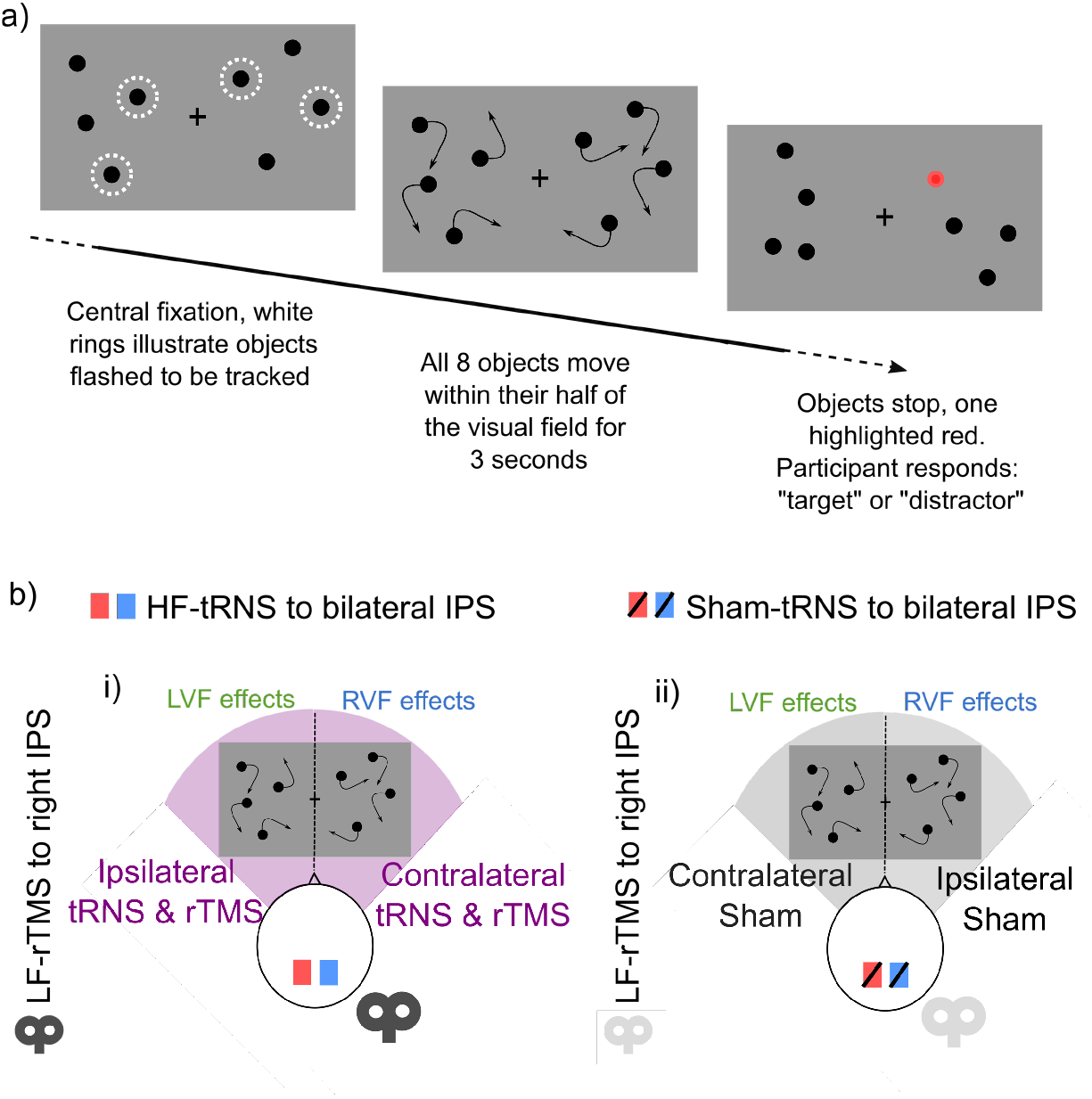
Sustained attention task & multi-stimulation montage for experiment 1. 1a) Bilateral multiple object tracking (MOT) was used to record participants sustained attention across time after stimulation. 1b) Depiction of each stimulation montage for experiment 1.

#### Thresholding stimuli

Participants performed bilateral MOT in a staircase procedure to establish each participants’ 75% correct threshold (Cornsweet, 1962). Participants first practiced MOT for 16 trials when the dots moved at the slow speed of 2 deg/sec. After the practice block, participants’ threshold was determined by changing the speed of the moving dots on each trial. Participants completed eight interleaved 3/1 staircases to assess their individual speed thresholds. The staircases increased the speed after three correct trials and reduced it following a single incorrect response. The staircases terminated after a combined 16 reversals, with threshold parameters estimated from the last 3 reversals. Speed was adjusted to yield 75% accuracy in the target/distractor judgments. Participants then performed bilateral MOT at their 75% correct threshold speed (degrees per second) for the duration of the experiment. In order to determine a behavioral change with stimulation, all participants should perform below ceiling at the same baseline prior to stimulation. Our participants 75% correct speed ranged from 5 –14.5 deg/sec for experiment 1 (Figure 2a).

**Figure 2:**
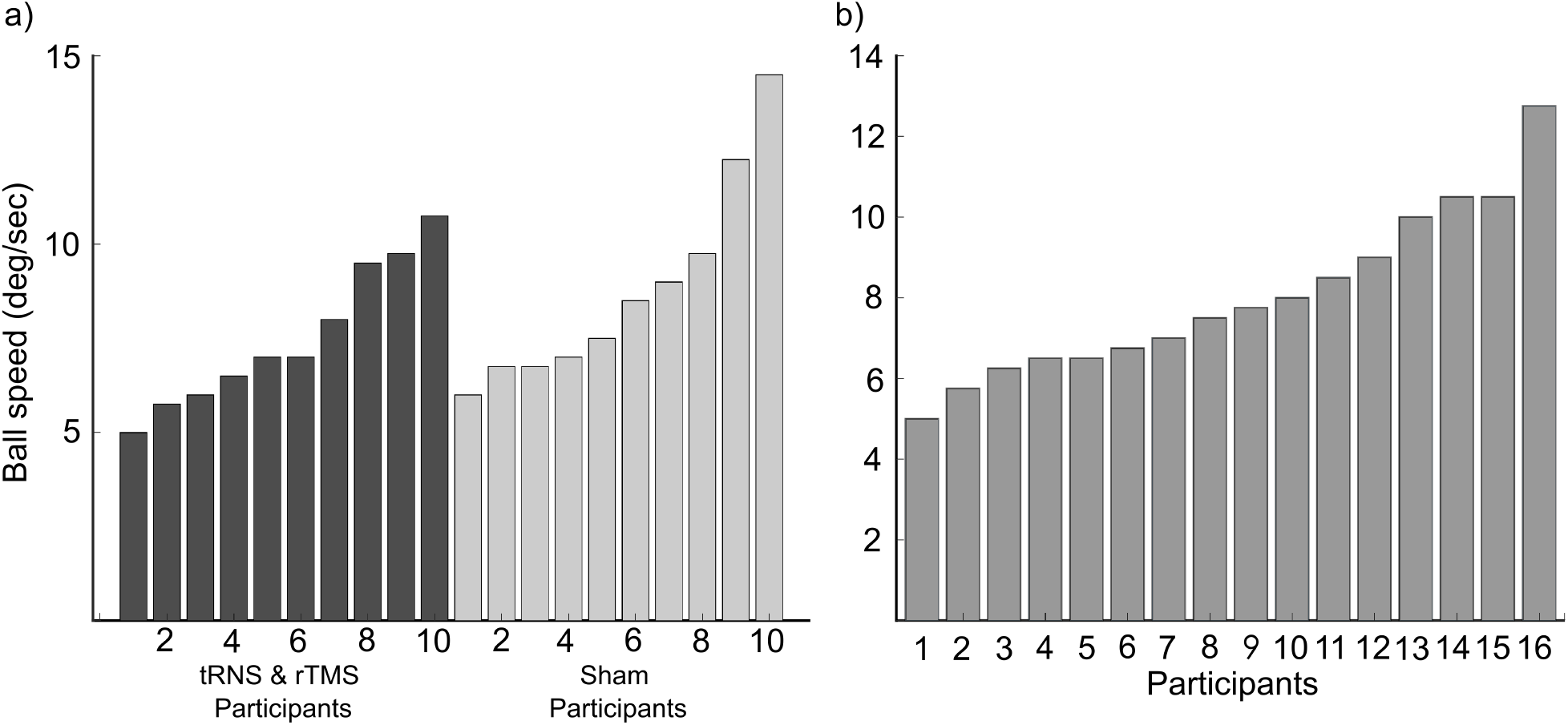
Threshold speed for each participant. 75% correct speed threshold in the bilateral MOT task was calculated for each participants using a staircase procedure. Data sorted lowest to highest for each group of participants. The speeds range from 5 – 14.5 degrees per second across experiments 1 (a) and 2 (b).

#### Experimental Designs & Statistical Analyses

In the experimental session, each participant was first tested on the bilateral MOT using a staircase procedure to determine the 75% correct speed threshold. This speed was kept fixed for the remainder of the experiment. Prior to stimulation, participants underwent six minutes of bilateral MOT to ensure equal performance across groups before stimulation. Each participant was randomly assigned to one of two stimulation protocols (Figure 1b: i) 20 minutes of high-frequency transcranial random noise stimulation (HF-tRNS) to bilateral posterior IPS followed by 15 minutes of low-frequency repetitive transcranial magnetic stimulation (LF-rTMS) to right IPS, ii) 20 minutes sham HF-tRNS to bilateral IPS followed by 15 minutes sham rTMS to right IPS. None of the participants had experienced brain stimulation previously, and therefore were unable to determine if they received real or sham stimulation. Participants also performed bilateral MOT during HF-tRNS stimulation. After stimulation, participants performed 94 minutes of MOT across 12 blocks of 6 minutes each, with two minutes break between each block. Each block comprised a total of 28 trials, with 19 trials with targets probed in the left (and 19 in the right) visual field. The order of the tested visual field was randomized across trials. Data were collected in 12 six-minute bins, however we collapsed the data every two bins to increase the number of trials per bin for analysis. This resulted in six 14-minute bins, 12-minutes plus the 2-minute break between the bins.

We fit a general linear mixed effects regression models to the object tracking accuracy using R (R Core Team, 2019) and the *glmer()* function within the *lme4* package (Bates et al., 2015). The interaction of interest was between stimulation and time, however we were also interested in visual field specific stimulation impact. Stimulation was the between-subjects predictor (*Sham-tRNS & rTMS* or *HF-tRNS & rTMS*), Time (6 bins: *0-14*, *16-30*, *32-46*, *48-62*, *64-78*, & *80-94*) and Visual Field (*Left* & *Right*) were the within-subject predictors. All these predictors were treated as categorical factors within the model, and random intercepts were included for participants. *Chi-squared* and *p-values* were reported for the interactions in the model and all the main effects. Model comparisons were also conducted to determine which model best predicted the data. We further present individual contrast between stimulation conditions at each time-point, controlled for multiple comparisons using *emmeans()* and *adjust = “mvt”* (Lenth et al., 2020).

To ensure there was no difference between groups prior to stimulation, we also performed a contrast between pre-stimulation behavior of the sham group and the group which received multi-method stimulation, by visual field.

Finally, we also performed a general linear mixed effect model on the behavior during tRNS stimulation for the sham and multimethod stimulation group across time. tRNS was performed for 20 minutes, so we blocked the behavior by first and last ten minutes to determine if there was a time-specific impact. Random intercepts were included for participants. *Chi-squared* and *p-values* were reported for the interaction model and all the main effects.

### Experiment 2: Multimethod stimulation in spatially specific reduction of attention deterioration

Experiment 2 was a within-subjects design with five independent sessions. Participants attended an fMRI session first to localize the stimulation sites, and then attended four separate sessions of non-invasive brain stimulation followed by behavioral testing. Each stimulation session involved a different stimulation protocol and was separated by at least 48 hours to avoid for stimulation carryover effects (Figure 3a).

**Figure 3:**
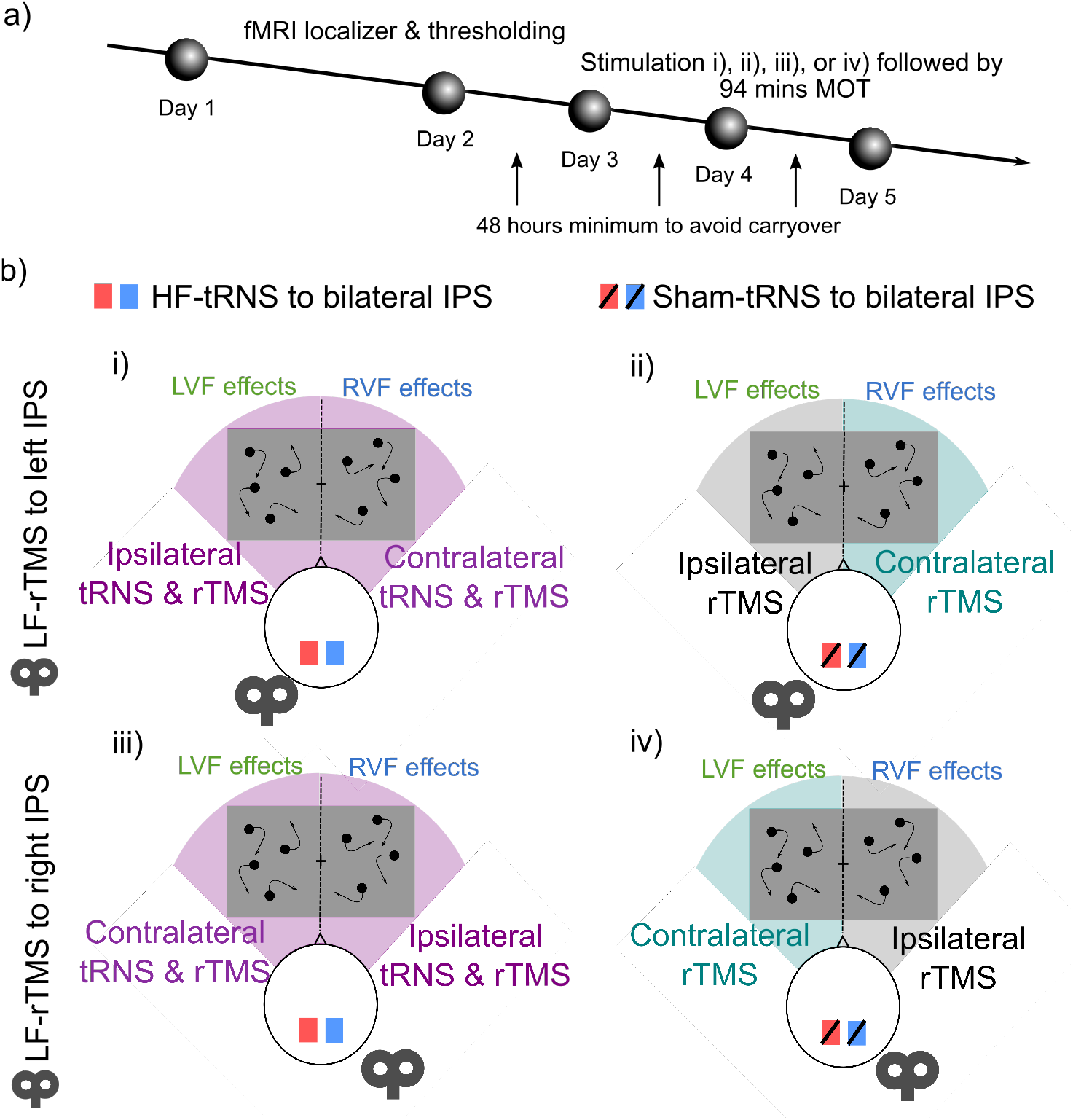
Multi-stimulation montage & protocol for experiment 2. a) Portrayal of five-day protocol. b) Depiction of each stimulation session for experiment 2. Note all left visual field stimulation effects were compared with left visual field ipsilateral rTMS control condition in a) ii), and right stimulation effects with right visual field ipsilateral rTMS control condition in a) iv).

#### Stimuli

During the localizer runs in the fMRI, participants viewed the stimulus on an fMRI compatible screen positioned in the bore of the magnet at a distance of 104 cm. The screen projected the stimulus from a 13-inch MacBook Pro (Retina; screen resolution: 1280 × 800). During behavioral testing after stimulation, the MOT was presented exactly the same as in experiment 1, where participants viewed the stimulus on the same 13-inch MacBook Pro at a distance of 60 cm.

#### Standard bilateral Multiple Object Tracking Paradigm (MOT)

Performed in the exact format as that of experiment 1.

#### Thresholding stimuli

Performed at the beginning of the session for each participant in the same staircase procedure as described for experiment 1. Participants’ 75% correct speed ranged from 5 – 12.75 deg/sec for experiment 2 (Figure 2b).

#### Localizer stimuli

Unilateral MOT was used to localize the posterior portion of the intraparietal sulcus (IPS) in both the right and left cortex. Posterior IPS is typically localized using fMRI with covert attention to the right and left visual field separately (Gitelman et al., 1999). During the localizer scan in the MRI, participants performed unilateral MOT for one run of 8.5 minutes. Each trial was performed similarly to the standard MOT described above, but two objects were cued to be tracked from one visual field only, rather than two objects in each visual field. Therefore, fixation remained central and attention was diverted to one visual field in each trial.

#### Experimental Designs & Statistical Analyses

The experiment was performed as a within-subjects design across five separate sessions, each participant performed all conditions of the experiment. On day one, participants performed unilateral MOT in the fMRI to localize the IPS for the following stimulation session, and subsequently perform the bilateral MOT using the staircase procedure to determine individual threshold speed at which each subject performed the task at 75% correct. Day two to day five were counterbalanced across subjects using Balanced Latin Squares. On each day the participant received one of four stimulation protocols (Figure 3b): 20 minutes of (i) high-frequency transcranial random noise stimulation (HF-tRNS) or (ii) sham tRNS to bilateral posterior IPS, followed by 15 minutes of low-frequency repetitive transcranial magnetic stimulation (LF-rTMS) to *left* IPS; or 20 minutes of (iii) HF-tRNS or (iv) sham tRNS to bilateral posterior IPS, followed by 15 minutes of LF-rTMS to right IPS.

After stimulation, the participant performed bilateral MOT for 94 minutes, twelve blocks of six minutes separated by two minutes of rest between each block (see procedure of *Experiment 1* for details).

We fit general linear mixed effects models to examine visual field specific impact of stimulation across time. As with experiment 1, we used the *lme4()* package in R to perform glmer (R core team, 2019; Bates et al., 2015). Our within-subjects predictors were Stimulation (*Contra-tRNS-rTMS, Ipsi-tRNS-rTMS, Contra-rTMS, & Ipsi-rTMS*), Time (6 bins: *0-14, 16-30, 32-46, 48-62, 64-78, & 80-94 minutes*), and Visual Field (*Left & Right*). Random intercepts were included for participants. We reported interactions and main effects with *Chi-squared* and *p-values*. Similar to experiment 1, we also performed model comparisons to ensure we had the model which best fit our data. We further presented individual contrasts between stimulation conditions at each time-point, controlled for multiple comparisons (using *emmeans()* & *adjust = “mvt”* in *contrast* (Lenth et al., 2020). Stimulation input for the generalized linear mixed effects model was considered by contralateral and ipsilateral visual field impact due to the active control condition, which was visual field specific. Evidence suggests attention performance differences between left and right visual field (Alvarez & Cavanagh, 2005; Chen et al., 2013), therefore a visual field specific control for each stimulation condition was necessary. In the LF-rTMS only sessions, object tracking performance in the hemifield ipsilateral to the LF-rTMS acted as the visual field specific active control. For instance, behavior in the left visual field during LF-rTMS to left IPS was the control behavior for the behavior in the left visual field in each of the other stimulation sessions. For an effective active control, a stimulation site is chosen as it is hypothesized to not impact the behavior of interest (Robertson et al., 2003; Valero-Cabré et al., 2017). In our case, rTMS to a region specifically localized for contralateral visual field attention was not expected to impact ipsilateral visual field attention (Dambeck et al., 2006; Battelli et al., 2009; Edwards et al., 2017).

#### MRI acquisition

In experiment 2, functional and anatomical MRI data were acquired using a 32-channel phased-array head coil with a 3 Tesla MRI system (Siemens Prisma) at the Harvard Center for Brain Sciences. For the functional scans, contrasts of blood oxygenation level-dependent (BOLD) activity were obtained using an echo-planar imaging sequence (parameters: 65 slices; slice thickness = 2.40 mm; FOV = 211 mm; 2.4 × 2.4 × 2.4 voxel size; Flip Angle = 64; TR = 1; TE = 32.60 ms; multi-band accelerator factor of 5). High-resolution T1 scans were acquired using 3D MPRAGE protocol (parameters: 176 slices; FOV = 256 mm; 1×1×1 mm voxel resolution; gap thickness = 0 mm; TR = 2200 ms; TE = 1.69 ms).

#### MRI analysis & localization of posterior IPS

The functional and anatomical data from experiment 1 were analyzed using BrainVoyager QX®. The first two volumes of each functional run were discarded to avoid saturation effects. Low-frequency noise and drift was removed using high-pass filtering during 3D-motion correction. The functional data was then aligned to the high-resolution anatomical data in native space. We convolved event timing with a hemodynamic model to generate predicted brain responses, with the associated beta weights estimated using a general linear model. To localize left and right posterior IPS, we contrasted the beta estimates for the unilateral MOT attend right > unilateral MOT attend left. With this contrast, activity in left IPS indicates lateralized attention to the right visual field, and activity in right IPS indicates lateralized attention to the left visual field (Figure 4). The most activated voxel within left and right posterior IPS was selected as the region for stimulation, individually measured for each subject (averaged coordinates across all subjects: Left IPS: x=-21.33(4.83), y=-83.53(3.01), z=19.00(4.87); Right IPS x=20.54(4.80), y=-83.35(4.19), z=17.96(5.56); depicted by red spheres in Figure 4). This data was saved in native space and retained for tRNS and TMS coil positioning.

**Figure 4:**
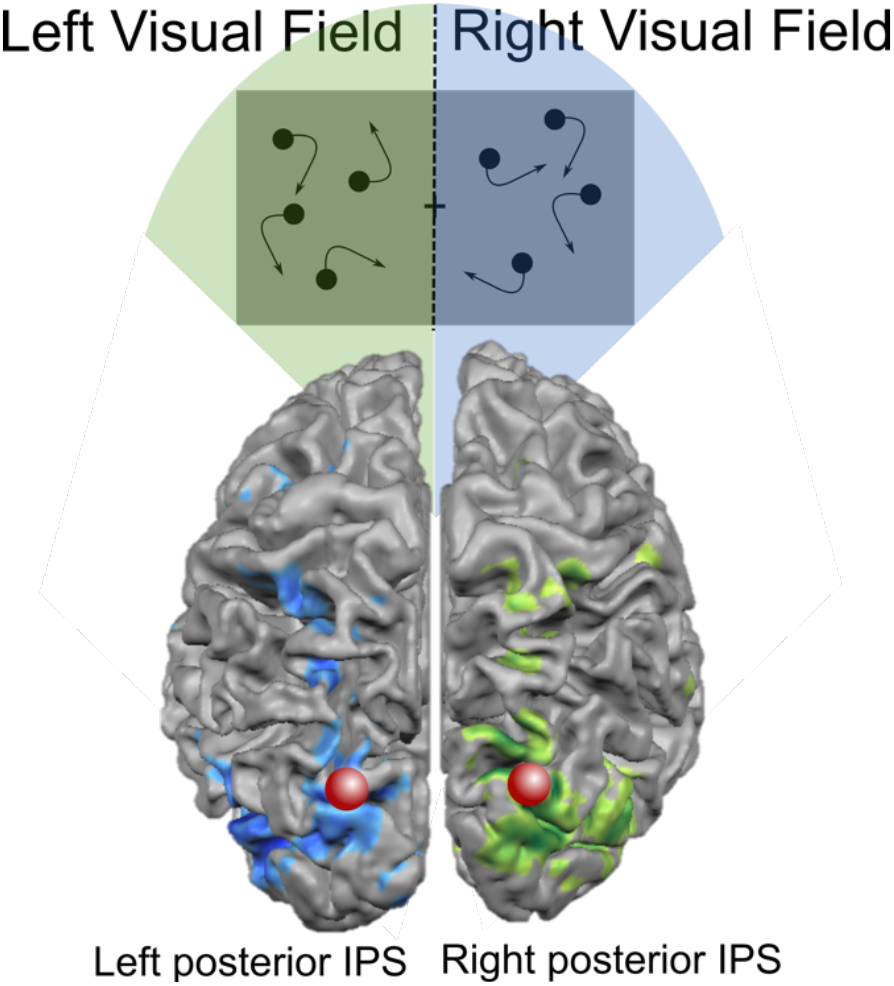
fMRI localizer for posterior bilateral intraparietal sulcus. Participants were presented with unilateral MOT in a fast-event related design. In each trial the participant had to fixate centrally and attend to targets in either the left or the right visual field, never both visual fields simultaneously. Right posterior IPS was localized through unilateral tracking in the left visual field, and left posterior IPS localized with unilateral tracking in the right visual field. The red spheres depict the average coordinates across all participants from experiment 1.

#### Stimulation Equipment & Coil Positioning

Stimulation sites for both tRNS and rTMS were selected based on fMRI data which localized each participants’ left and right posterior IPS (see *MRI Analysis & localization of posterior IPS* and Figure 4) in experiment 2 and averaged brain coordinates for experiment 1. In experiment 2, Brainsight Frameless Stereotaxy System (Rogue Research, Montreal, Canada) was used to align the participant with their native space functional MRI data. In experiment 1, the averaged coordinates from experiment 2 were used to guide stimulation target. Participants were aligned to the model brain within Brainsight Frameless Stereotaxy System (Rogue Research, Montreal, Canada), the model brain size was adjusted to each individual participant, and then the averaged coordinates were selected for target. HF-tRNS was delivered for 20 minutes using electrodes and a DC-Stimulator (Eldith-Plus, Neuroconn), at 1mA at random frequencies between 101-640 Hz. Stimulation began and finished with a 15-second fade-in/ fade-out ramp. The electrodes were placed inside saline soaked sponges, and were positioned over left and right IPS. tRNS stimulation delivered using this protocol has been shown to have an excitatory impact on the cortex (Herpich et al., 2018). The procedure for Sham-tRNS was exactly the same, with a fade-in and fade-out ramp, but stimulation was turned off and not delivered during the 20 minutes. LF-rTMS was delivered using a MagPro® by MagVenture with a figure-8 coil with an inbuilt cooling system. Stimulation was performed at 1Hz with 65% machine output intensity (Battelli et al., 2009). The coil was held with the handle pointing backward, in a tangential orientation over the either left or right posterior IPS (Ruff et al., 2008; Battelli et al., 2009; Edwards et al., 2017). For the sham rTMS condition in experiment 1, the coil was flipped away from the cortex, but in the same tangential orientation over the right posterior IPS. The participants experienced the set-up and auditory stimulation of rTMS, without any magnetic stimulation. Anecdotally, on average participants were unable to determine if they had received sham in experiments 1 or 2 at the end of the study.

## Results

### Experiment 1: Multimethod stimulation to prevent attention deterioration

We examined the effect of multi-method non-invasive brain stimulation on bilateral attention, dependent of visual field. After HF-tRNS to bilateral IPS, followed by LF-rTMS to IPS, we expected to prevent attention deterioration across time.

#### Pre-stimulation performance

Due to the between-subjects design, we ran a post thresholding, pre-stimulation block of trials to confirm there were no differences between the sham and stimulation groups. There was no significant pre-stimulation difference between tRNS & rTMS and sham by visual field (χ^2^(1)=0.082, p=0.775), and no main effects of stimulation nor visual field (p>0.05). This indicates the two groups were not significantly different prior to stimulation.

#### During stimulation performance

We also examined the impact of tRNS on behavior during stimulation. We found no interaction between stimulation, time and visual field (χ^2^(1)=0.506, p=0.477; Figure S1), or any two way interactions, nor main effects (p>0.05). Therefore, tRNS did not significantly impact behavior during stimulation in comparison to sham.

#### Post-stimulation performance

We analyzed the impact of multimethod stimulation across 94 minutes in comparison to sham and determined if there is a visual field specific effect.

We found a main effect of time on performance (χ^2^(5)=66.4683, p<0.0001, *glmer*), but no main effect of stimulation (χ^2^(1)=1.1685, p=0.2797, *glmer)* or visual field χ^2^(5)=1.9147, p=0.1664, *glmer).* When examining the interactions, we found no significant interaction between stimulation, time, and visual field (χ^2^(5)=3.8312, p=0.5739, *glmer*). However, we did find a significant interaction between stimulation and time (χ^2^(5)=35.6657, p<0.0001, *glmer*; Figure 5a). This indicated that there was a difference between sham and multi-method stimulation across time, but this difference was not modulated by visual field (Figure 5b & c). Model comparison analysis also supported stimulation × time interaction as the best fit for the data (see Supporting Information). Therefore, time from stimulation offset significantly impacted the change in behavior. When examining individual time-points to determine when there was a significant difference in attention following stimulation, we found performance was significantly lower in participants for the sham group in comparison to the stimulation group at 64-78 minutes (estimate=1.475, se=0.589, z=2.505, p=0.0497) and at 80-94 minutes post-stimulation (estimate=0.630, se=0.235, z=2.686, p=0.0303; *emmeans* with *adjust “mvt”*).

**Figure 5:**
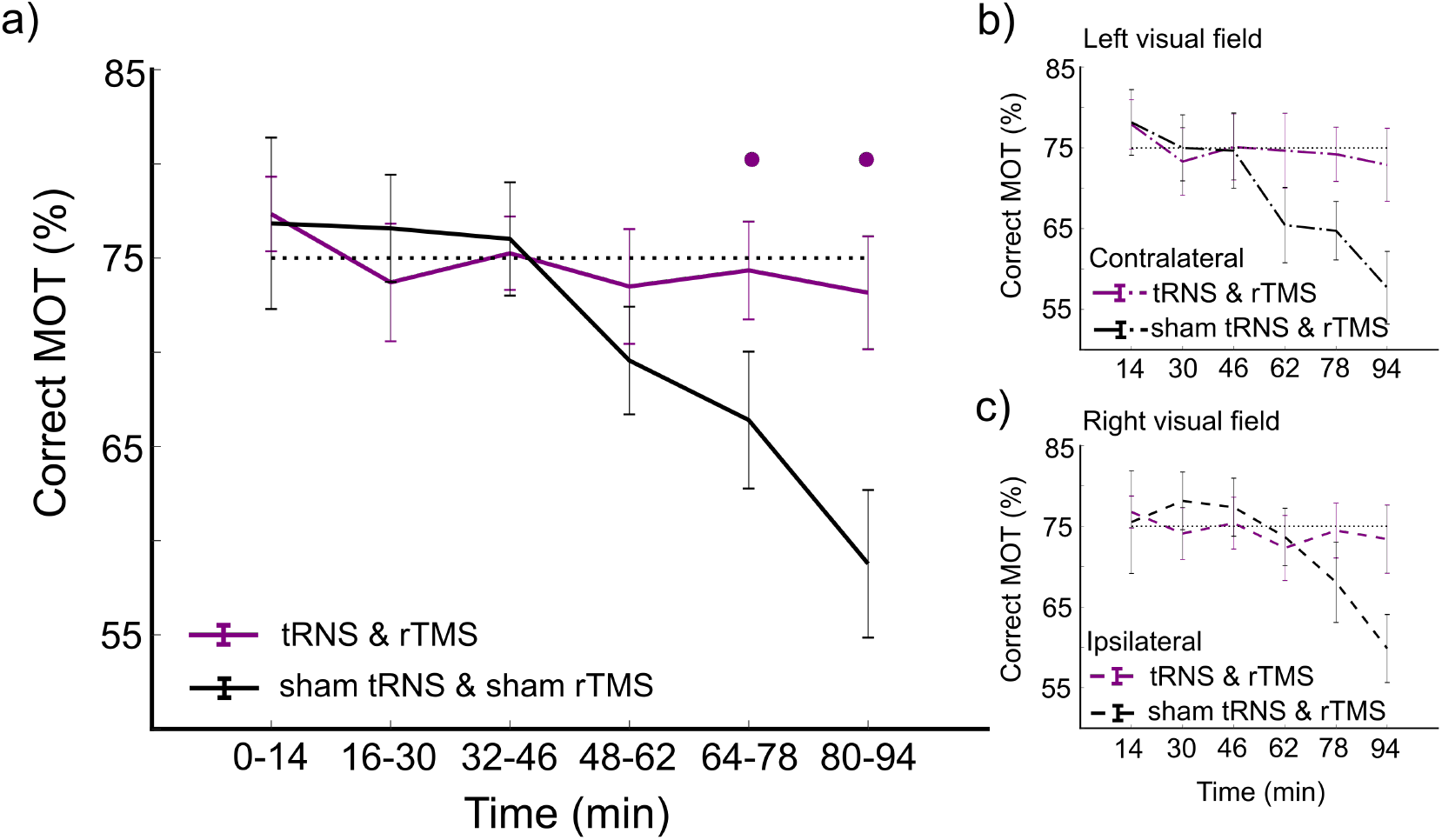
Experiment 1: Stimulation impact on MOT performance. a) Impact of tRNS & rTMS on MOT collapsed across visual field. b) Left visual field stimulation effects: Stimulation impact on MOT contralateral to tRNS & rTMS relative to sham. c) Right visual field stimulation effects: Stimulation impact on MOT ipsilateral to tRNS & rTMS relative to sham. • indicates p<0.05 adjusted for multiple comparisons.

### Experiment 2: *Multimethod stimulation in spatially specific reduction of attention deterioration*

In Experiment 2 we expected to reproduce the lack of attention deterioration up to 94 minutes after stimulation with tRNS priming prior to LF-rTMS. Furthermore, we performed this experiment to determine if targeting left or right IPS with multi-method stimulation differentially modulated sustained attention in the left and right visual fields.

Importantly, we added another stimulation condition to determine if rTMS alone could result in prevention of attention deterioration up to 94-minutes. rTMS over IPS has been shown to have contralateral visual field specific impact (Dambeck et al., 2006; Battelli et al., 2009). Therefore, in the rTMS alone stimulation day, we assume the behavior in the contralateral visual field as impacted by rTMS, whereas the ipsilateral behavior acts as an active control, specific to visual field.

#### Active control site: Ipsilateral visual field from rTMS

First, we determined if behavior in the ipsilateral visual field from rTMS was a strong control for behavior. We found the ipsilateral visual field was not significantly different from 75% for all six time-points (p>0.05) in both the left and right visual field (Figure 6a & b lower inlay). Each participant was thresholded to 75% correct prior to intervention, therefore no deviation from 75% indicates no change in behavior due to rTMS in the ipsilateral visual field.

**Figure 6:**
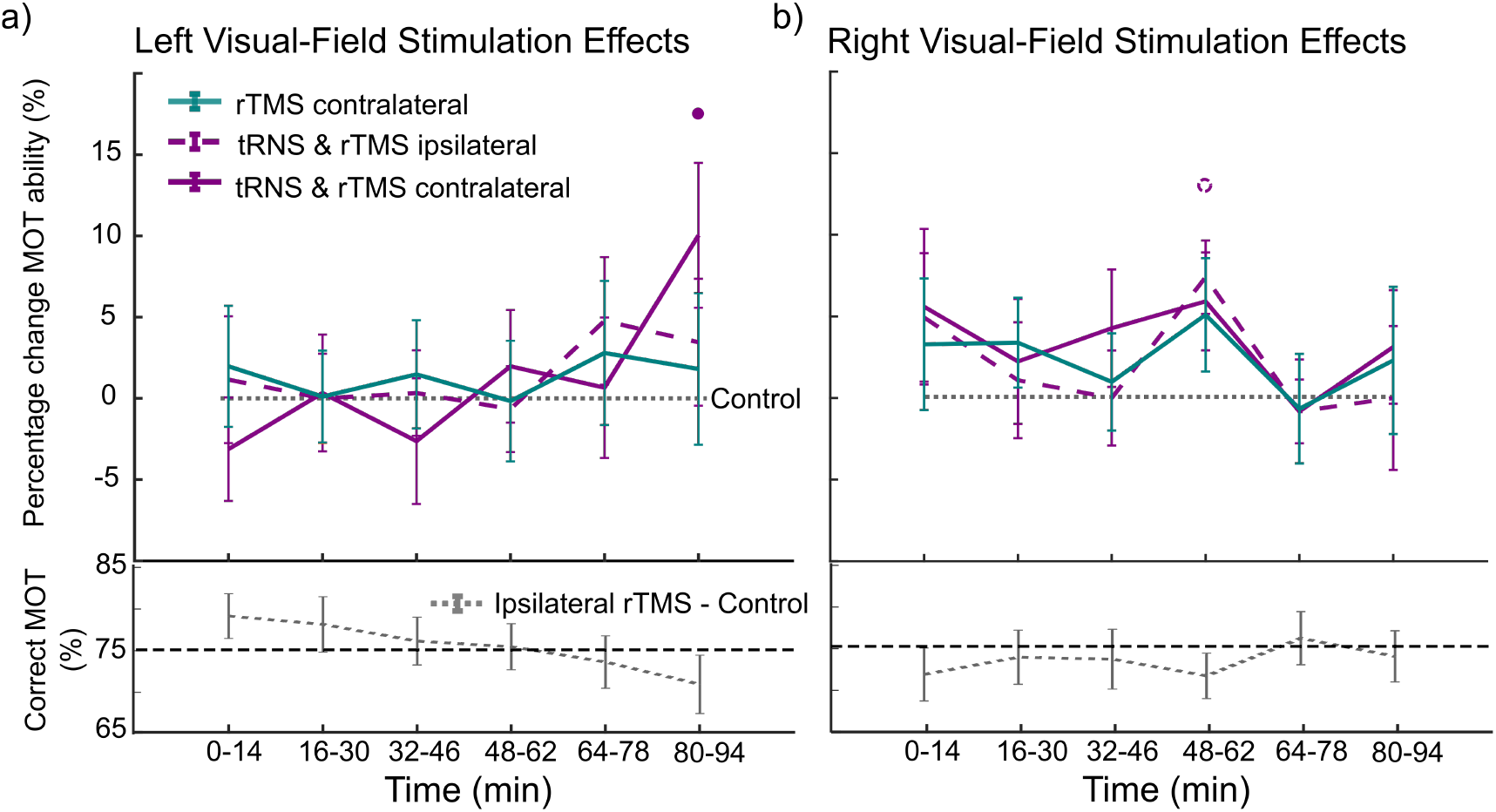
Experiment 2: Stimulation impact on MOT performance in each visual field across time. a) Stimulation impact on MOT in the left visual field. b) Stimulation impact on MOT in the right visual field. The grey dotted line at the bottom of the image depicts the percentage control MOT performance across time. The percentage change in performance after each stimulation condition is then presented above. The • indicates p<0.05; the dotted • indicates significant prior to multiple comparisons.

Surprisingly, no change in attention across time in our control condition indicated that the participants from experiment 2 did not experience sustained attention decrement. We believe the lack of sustained attention decrement was due to possible training effects across the four sessions of the experiment. Testing this hypothesis, we found a significant impact of session (χ^2^(3)=9.6934, p=0.02136) on accuracy in MOT, with performance at 80-94 minutes in session 3 and session 4 significantly better than in session 1 (Session 4 vs. Session 1: estimate = 0.522, se = 0.189, z-ratio = 2.765, p = 0.0159; Session 3 vs. Session 1: estimate = 0.477, se = 0.188, z-ratio = 2.536, p =0.0310; Figure S2). This stimulation-type independent increase in task performance across session could account for the lack of sustained attention decrement in control conditions. Through counterbalancing the order of stimulation protocols across sessions, a subset of subjects had their control sessions last, introducing variability in performance, and increasing baseline sustained attention. Irrespective of the lack of decrement, we analyzed the multi-method stimulation effect to determine if it still caused a boost in attention in comparison to the control behavior.

#### Impact of stimulation on sustained attention

Percentage correct MOT is reported in Figure 6 with ipsilateral rTMS control condition in the bottom inlay, and percentage change for each stimulation condition in comparison to the control, reported above. Using generalized linear mixed effects modelling, we found a main effect of stimulation, (χ^2^(3)=11.3708, p=0.009) and visual field (χ^2^(1)=8.8315, p=0.002), but no effect of time (χ^2^(5)=10.0352, p=0.074). We also found a significant three-way interaction of stimulation × time × visual field (χ^2^(15)=26.9615, p=0.029). The stimulation × time × visual field interaction was further supported as the best fit for the data according to model comparisons (see Supporting Information). This indicates that attention was modulated by stimulation across time and this modulation was dependent upon visual field (Figure 6).

When computing contrasts on estimated marginal means, we found a significant difference in tracking performance at 80-94 minutes between the left visual field contralateral to tRNS & rTMS to right IPS and control (estimate = 0.572, se = 0.138, z-ratio = 4.157, p=0.0006; *emmeans(),* with *adjust “mvt”*). In the right visual field (ipsilateral to tRNS & rTMS to right IPS), there was also a trend in sustained attention increase, however not to significance (estimate = 0.395, se = 0.135, z-ratio = 2.917, p=0.0584; controlled for multiple comparisons). No other time-point was significantly different from control.

This analysis indicates that right hemisphere multi-method stimulation impacts predominately contralateral left visual field sustained attention, also with a trend to impact ipsilateral right visual field attention. Whereas left hemisphere multi-method stimulation does not seem to impact contralateral or ipsilateral attention.

rTMS alone also does not seem to have an impact on attention at any time-point, regardless of stimulation site (p<0.05).

## Discussion

We investigated the use of multi-method brain stimulation as an intervention to improve sustained attention over time. In two experiments, we found multi-method stimulation maintained and improved sustained attention up to 94 minutes post-stimulation. This is the first evidence to suggest multi-stimulation has a lasting impact on cognitive behavior, complementing the previous studies demonstrating lasting impact to physiological response (Iyer et al., 2003; Siebner et al., 2004; Lang et al., 2004; Fricke et al., 2010; Bocci et al., 2014).

In previous experiments, high-frequency stimulation immediately followed by low-frequency stimulation has resulted in lasting inhibition, demonstrated physiologically by decreased motor evoked potential amplitude (Iyer et al., 2003), and visual evoked potential amplitute (Bocci et al., 2014). Our effect shows multi-method stimulation improved sustained attention, seemingly at odds with the previous inhibitory multi-method physiological responses. It is important to restate that our effect is likely due to a dynamic change of excitatory-inhibitory balance across the whole dorsal attention network. Like previous multi-method experiments, we expected multi-method stimulation to inhibit IPS, but unlike previous multi-method experiments, we expected the inhibition to cause compensatory activity from other nodes of the dorsal attention network (Paus et al., 1997; Grefkes et al., 2009; Lee & D’Esposito, 2012; Plow et al., 2014; Battelli et al., 2017). Experiments recording the impact of a single method protocol of LF-rTMS to the IPS have shown a late boost in attention capability in patients (Brighina et al., 2003; Agosta et al., 2014) and a functional reorganization of the attention network beginning 36 minutes and lasting for 50 minutes after stimulation in healthy participants (Battelli et al., 2017). We believe that the lack of attention deterioration following multi-method stimulation, is a result of functional reorganization, like that of Battelli et al., (2017). Although behavioral change was not previously detected with network-wide function change following LF-rTMS alone (Battelli et al., 2017), multi-method stimulation may have caused a sustained enough inhibitory effect to result in behavioral impact. The lateness of the behavioral impact is likely due to stimulation improving sustained attention decrement, which begins to deteriorate within 75 minutes (Whitehurst et al., 2019).

In line with our hypothesis, our first experiment demonstrated a significant impact of tRNS & rTMS on improving sustained attention, regardless of visual field. This indicates that compensation following multimethod stimulation to right IPS impacted attention in both visual fields equally. Our second experiment further demonstrated this effect was specific to multimethod stimulation to right IPS. Multimethod stimulation to right IPS significantly impacted attention in the left visual field, with a trend of an increase in attention in the right visual field. However, we found no impact of multimethod stimulation over the left parietal cortex in either the right or left visual field. Bilateral visual field impact from multi-method stimulation targeted at the right parietal cortex supports the evidence of bilateral visual field representation in the right parietal cortex (Sheremata et al., 2010: Sheremata & Silver 2015). The visual field wide attention increase following multi-method stimulation to right IPS may be useful in designing new strategies to sustain and improve attention in healthy and neurological populations.

Along with a late boost in attention, one might expect an initial decrease in attentional capability following HF-tRNS and LF-rTMS multi-method stimulation. The initial decrement in attention could demonstrate the magnified inhibition expected directly following stimulation, however a decrease in attention capability was not found. The lack of behavioral decrement could be explained by the interaction of posterior IPS and its homotopic counterpart in the other hemisphere following targeted inhibitory brain stimulation. Using a different, but mechanistically similar, method to inhibit cortical function, researchers have found that an eye deprived of visual stimuli (through eye patching) *strengthens* in perceptual response directly after deprivation (Mrsic-Flogel et al., 2007; Lunghi et al., 2015; Kim, Kim and Blake 2017). Long-term potentiation (LTP) following deprivation and non-invasive brain stimulation hinge on similar underlying mechanisms including activation of NMDA receptors, concomitant GABAergic inhibition, and the production of acetylcholine neurotransmitter (Boroojerdi et al., 2001; Cheeran et al., 2009). This strengthening following monocular deprivation is in contrast to the usual functional decline associated with cortical inhibition. Even a short deprivation of 15 minutes can produce temporary strengthening of the deprived eye resulting in a change in perceptual behavior (Kim, Kim & Blake 2017). Similar to the mutual inhibition found between visual field specific attentional processing regions in temporoparietal areas, bistable perception between the two eyes is controlled via mutual inhibition between the eyes in early visual areas (Binda et al., 2018). In monocular deprivation, a boost in performance of the deprived eye was thought to be due to the subsequent lack of inhibition from the functioning eye (Lunghi et al., 2011). This lack of inhibition is demonstrated by the overall decrease in GABA concentration in the early visual cortex which correlates with post-deprivation perceptual performance of the deprived eye (Lunghi et al., 2015). Therefore, the boost in functionality may be a method for stabilizing homeostatic gain response, where the deprived cortex attempts to restore balance between the homotopic brain regions (such as binocular balance, Zhou, Clavagnier, & Hess, 2013). This theory could indeed account for the effects we found following multi-stimulation of the posterior IPS. Posterior IPS is reliant on mutual inhibition with its homotopic counterpart to adequately induce lateralized attention (Corbetta et al., 2005). It may be that the multi-method stimulation still has an inhibitory effect, but this is mitigated in the behavior by the reduction of mutual inhibition from homotopic posterior IPS. Together these effects could cancel one-another, resulting in no change in behavior.

Interestingly, there was also no effect of LF-rTMS alone to the posterior IPS directly after stimulation, further supporting our interpretation of the initial null effect in the multi-method condition. LF-rTMS can result in an acute inhibitory behavioral response directly after stimulation (e.g. Battelli et al., 2009; Edwards et al., 2017), but has also been reported to have no immediate but a *delayed* effect after stimulation to posterior IPS (Plow et al., 2014; Agosta et al., 2014), and occasionally, no impact on behavior at all (Edwards et al., 2017).

HF-tRNS stimulation seemed to have no direct impact on behavior during stimulation in experiment 1. Although cortical excitability has been demonstrated almost instantaneously following 20-minutes HF-tRNS (Snowball et al., 2013; Herpich et al., 2018), the excitability impact can take time to present during stimulation (Tyler et al., 2018) and it can show a cumulative effect across days and sessions (Herpich et al., 2019). Tyler et al., (2018) illustrated the cortical excitability profile during one session of HF-tRNS to bilateral IPS, with attention increasing significantly above control 25 minutes following stimulation. In experiments 1 and 2, it is therefore likely that cortical excitability following HF-tRNS had built significantly in bilateral IPS by the time LF-rTMS was applied.

Finally, regardless of the order of stimulation protocols used in experiment 2, participants’ sustained attention did not deviate from the performance threshold set at the beginning of the experiment. This lack of depreciation of attention across time indicates a training effects across session, further supported by our analysis. Our data is in line with previous studies which demonstrate multiple sessions of training can improve cognitive performance (Jaggi et al., 2008; Herpich et al., 2019; Pergher et al., 2019). Despite the impact of training across session, we still found a significant improvement in attention within the tRNS & rTMS session.

## Conclusion

Sustained attention is a limited resource, necessary across multiple cognitive tasks (DeGangi & Porges, 1990). Maintenance of attention across time is sought after in both the clinical and healthy populations. Here, we demonstrate attention can be maintained without decrement for up to (and potentially beyond) 94 minutes following multi-method brain stimulation. Future research should be focused on the underlying network changes following multi-method stimulation. Network based compensation for inhibition of one focal node of a network using multi-method stimulation may prove useful in other cognitive domains, such as working memory (Compte et al., 2000). Efficiency of network-wide communication has been demonstrated to be of utmost importance in conserving cognitive reserve with age (Weiler et al., 2018).

## Acknowledgements

Data collection funded by the Harvard Mind Brain Behavior Interfaculty Initiative. We thank Emily D. Grossman for her helpful insights.

