## Supplemental Materials for "Controlling brain state prior to stimulation of parietal cortex prevents deterioration of sustained attention"

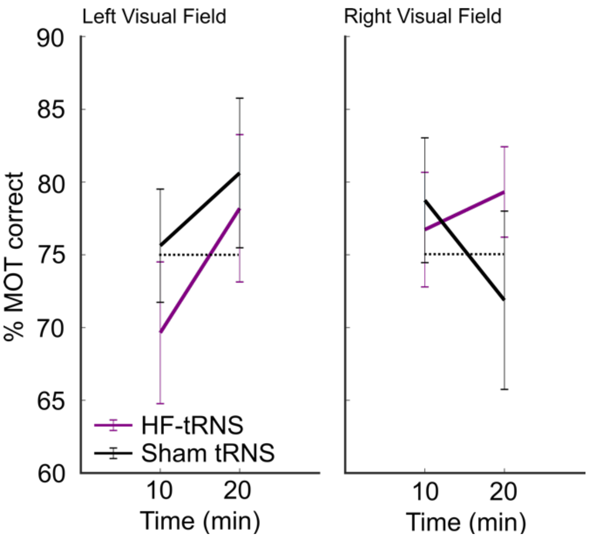


**Supplemental Figure 1: Percentage correct MOT performance during HF-tRNS and Sham-tRNS in Experiment 1.** Data split into left and right visual field, and first and last 10 minutes of stimulation.

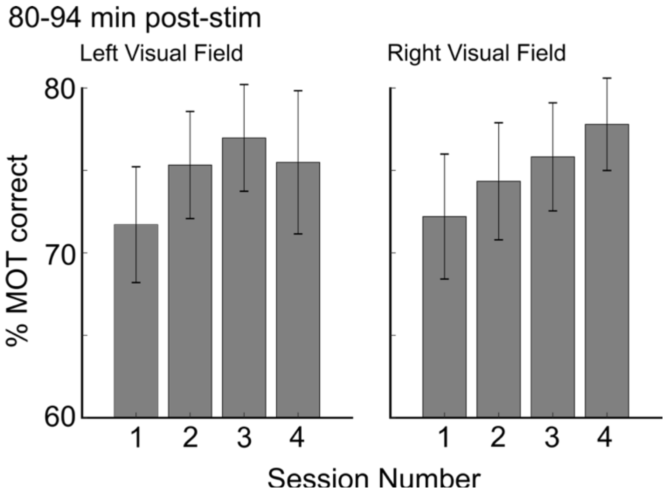


**Supplemental Figure 2: Impact of multiple session MOT on sustained attention in Experiment 2.** Percentage multiple object tracking accuracy change across session in the 80-94 minute time-bin post-stimulation.

### Model Comparisons

Model comparisons were conducted on various *glmer* models to determine the best model fit for the data collected from experiment 1 and experiment 2.

#### Model Comparisons – Experiment 1

**Syntax:**

First we compare our hypothesized model of best fit (Mod2 – interaction between Stimulation and Time) with a simpler model (Mod1 – no interaction):

```
Mod1 <- glmer (Accuracy ~ Stimulation + Times + VF + (1|Subs), data = Multimethod, family = binomial)
Mod2 <- glmer (Accuracy ~ Stimulation * Times + VF + (1|Subs), data = Multimethod, family = binomial)

anova(Mod1,Mod2)
```

| Mod | Df | AIC | BIC | loglik | deviance | Chisq | Chi Df | Pr(>Chisq) |
|---|---|---|---|---|---|---|---|---|
| Mod1 | 9 | 10411 | 10475 | -5196.5 | 10393 | | | |
| Mod2 | 14 | 10385 | 10485 | -5178.6 | 10357 | 35.866 | 5 | <0.000001 |

This demonstrates Mod2 is a better fit for our data than Mod1.

Next we add a level of complexity by determining if the additional variable of visual field in the interaction increases the fit of the model to our data:

```
Mod5 <- glmer (Accuracy ~ Stimulation * Times * VF + (1|Subs), data = Multimethod, family = binomial)

anova(Mod2,Mod5)
```

| Mod | Df | AIC | BIC | loglik | deviance | Chisq | Chi Df | Pr(>Chisq) |
|---|---|---|---|---|---|---|---|---|
| Mod2 | 14 | 10385 | 10485 | -5178.6 | 10357 | | | |
| Mod5 | 25 | 10398 | 10576 | -5173.9 | 10348 | 9.2764 | 11 | 0.5964 |

The lack of significant difference between Mod5 and Mod2 indicates that the additional complexity of Mod5 did not improve the fit to the data, therefore we reject Mod5 in favor of Mod2.

We also present Mod3 and Mod4 with interaction terms which also did not fit the data better than the simplest model, Mod1:

```
Mod3 <- glmer (Accuracy ~ Stimulation + Times * VF + (1|Subs), data = Multimethod, family = binomial)

anova(Mod1,Mod3)
```

| Mod | Df | AIC | BIC | loglik | deviance | Chisq | Chi Df | Pr(>Chisq) |
|---|---|---|---|---|---|---|---|---|
| Mod1 | 9 | 10411 | 10475 | -5196.5 | 10393 | | | |
| Mod3 | 14 | 10418 | 10518 | -5195.2 | 10390 | 2.6514 | 5 | 0.735 |

```
Mod4 <- glmer (Accuracy ~ Times + Stimulation * VF + (1|Subs), data = Multimethod, family = binomial)
```

anova(Mod1,Mod4)

| Mod | Df | AIC | BIC | loglik | deviance | Chisq | Chi Df | Pr(>Chisq) |
|---|---|---|---|---|---|---|---|---|
| Mod1 | 9 | 10411 | 10475 | -5196.5 | 10393 | | | |
| Mod4 | 10 | 10410 | 10481 | -5.195.1 | 10390 | 2.7434 | 1 | 0.09766 |

### Model Comparisons – Experiment 2

### Syntax:

We find the model which best fits our data is the model including the three way interaction between Stimulation x Time x Visual Field:

Mod1 <- glmer (Accuracy ~ Stimulation + Times + VF + (1|Subs), data = Multimethod, family = binomial)
Mod5 <- glmer (Accuracy ~ Stimulation * Times * VF + (1|Subs), data = Multimethod, family = binomial)

| Mod | Df | AIC | BIC | loglik | deviance | Chisq | Chi Df | Pr(>Chisq) |
|---|---|---|---|---|---|---|---|---|
| Mod1 | 11 | 31465 | 31556 | -15722 | 31443 | | | |
| Mod5 | 49 | 31485 | 31891 | -15694 | 31387 | 56.31 | 38 | 0.02817 |

No simpler model fit the data significantly:

Mod2 <- glmer (Accuracy ~ Stimulation * Times + VF + (1|Subs), data = Multimethod, family = binomial)

anova(Mod1,Mod2)

| Mod | Df | AIC | BIC | loglik | deviance | Chisq | Chi Df | Pr(>Chisq) |
|---|---|---|---|---|---|---|---|---|
| Mod1 | 11 | 31465 | 31556 | -15722 | 31443 | | | |
| Mod2 | 26 | 31478 | 31694 | -15713 | 31426 | 17.125 | 15 | 0.3115 |

Mod3 <- glmer (Accuracy ~ Stimulation + Times * VF + (1|Subs), data = Multimethod, family = binomial)

anova(Mod1,Mod3)

| Mod | Df | AIC | BIC | loglik | deviance | Chisq | Chi Df | Pr(>Chisq) |
|---|---|---|---|---|---|---|---|---|
| Mod1 | 11 | 31465 | 31556 | -15722 | 31443 | | | |
| Mod3 | 16 | 31465 | 31597 | -15716 | 31443 | 10.514 | 5 | 0.06192 |

Mod4 <- glmer (Accuracy ~ Times + Stimulation * VF + (1|Subs), data = Multimethod, family = binomial)

anova(Mod1,Mod4)

| Mod | Df | AIC | BIC | loglik | deviance | Chisq | Chi Df | Pr(>Chisq) |
|---|---|---|---|---|---|---|---|---|
| Mod1 | 11 | 31465 | 31556 | -15722 | 31443 | | | |
| Mod4 | 14 | 31469 | 31585 | -15721 | 31441 | 2.2816 | 3 | 0.516 |
